# Rapid hypermutation B cell trajectory recruits previously primed B cells upon third SARS-CoV-2 mRNA vaccination

**DOI:** 10.1101/2022.03.01.482462

**Authors:** Lisa Paschold, Bianca Klee, Cornelia Gottschick, Edith Willscher, Sophie Diexer, Christoph Schultheiß, Donjete Simnica, Daniel Sedding, Matthias Girndt, Michael Gekle, Rafael Mikolajczyk, Mascha Binder

## Abstract

High antibody affinity against the ancestral SARS-CoV-2 strain seems to be necessary (but not always sufficient) for the control of emerging immune-escape variants. Therefore, aiming at strong B cell somatic hypermutation - not only at high antibody titers - is a priority when utilizing vaccines that are not targeted at individual variants. Here, we developed a next-generation sequencing based SARS-CoV-2 B cell tracking protocol to rapidly determine the level of immunoglobulin somatic hypermutation at distinct points during the immunization period. The percentage of somatically hypermutated B cells in the SARS-CoV-2 specific repertoire was low after the primary vaccination series, evolved further over months and increased steeply after boosting. The third vaccination mobilized not only naïve, but also antigen-experienced B cell clones into further rapid somatic hypermutation trajectories indicating increased affinity. Together, the strongly mutated post-booster repertoires and antibodies deriving from this may explain why the booster, but not the primary vaccination series, offers some protection against immune-escape variants such as Omicron B.1.1.529.

**Brief summary:** Priming SARS-CoV-2 vaccinations generate antibodies from low-level matured B cells while the third vaccination strongly boosts somatic hypermutation potentially explaining different protection from immune-escape variants.

## Introduction

Until February 2022, the World Health Organization (WHO) counted 400 million severe acute coronavirus disease 2019 (COVID-19) infections caused by respiratory syndrome coronavirus 2 (SARS-CoV-2). By then, the number of deaths had totaled almost 6 million individuals globally. While mRNA-based and adenovirus-vectored vaccines have been developed at unprecedented speed, global vaccination strategies remain challenging and new SARS-CoV-2 variants with varying potential to evade adaptive immunity and/or to enhance transmissibility constantly emerge. Since memory B cell populations play a decisive role in severity reduction of COVID-19 and early antibody-mediated virus neutralization may even prevent infections, understanding infection- and vaccine-induced SARS-CoV-2 specific B cell immunity is critical (1–3).

As recently reviewed by Laidlaw et al. (4), COVID-19 generates both germinal center and extrafollicular B cell responses in unvaccinated individuals – depending on the severity of infection - that converge on B cells expressing antigen receptors with preferential immunoglobulin heavy chain variable-joining gene (IGHV-J) usage (1, 5–12). Interestingly, even B cells with low or absent IGHV affinity maturation can generate antibodies that specifically recognize and neutralize the ancestral strain of SARS-CoV-2 (6, 11, 13–15). Yet, a continued evolution of the humoral response appears to take place over at least six months after infection – even without re-infection – as demonstrated by sustained acquisition of IGHV somatic hypermutation despite waning antibody titers (16–21). There is emerging evidence that these rather mitigated B cell maturation dynamics may also be characteristic for vaccine-induced anti-SARS-CoV-2 immune responses (22, 23).

With the advent of SARS-CoV-2 variants of concern, affinity to the ancestral strain’s S protein (currently used in all licensed vaccines) does not necessarily predict antibody neutralization potency. Immune escape can affect clones with high affinity against the ancestral strain but, depending on the targeted epitope, some clones also retain their neutralizing potency against variants of concern (24). This is in clear contrast to clones with low affinity to ancestral SARS-CoV-2 that constantly fail to neutralize variants (24). This finding suggests that high affinity binding to the ancestral strain may provide some flexibility in compensating the effect of individual immune escape mutations (24). Therefore, in times of emerging viral variants an optimal vaccination strategy should aim at inducing the highest possible level of affinity maturation through somatic hypermutation, even if the available vaccines are targeted at the ancestral strain.

In the study presented here, we used two cohorts of not previously infected patients to compare antibody levels and somatic hypermutation trajectories across a primary series of two standard vaccinations with those induced by a third “booster” vaccination using immune repertoire sequencing. We show that B lineage evolution is low after the priming vaccinations. In contrast, the maturation trajectories induced by the third vaccination is compatible with selective mobilization and germinal center recruitment of naïve but also previously matured memory B cell lineages to undergo fast and extensive somatic hypermutation. This considerable affinity maturation and the resulting high antibody titers may explain the increased protection of the third “booster” vaccination against variants such as the Omicron variant B.1.1.529.

## Results

### Surveys, data collection and biobanking in NIS635

For NIS635, we used a classical biobanking study (HACO) and a digital cohort study (DigiHero) with flexible recruitment of participants into different survey modules to obtain COVID-19 and vaccination data in a large cohort and to acquire relevant biological samples from subgroups of interest. In DigiHero, 514 participants donated blood and completed the survey of the COVID-19 module until December 2021. 4,670 participants completed the survey on the third SARS-CoV-2 “booster” vaccination until the same data cut. In HACO, data and sample collection had been completed in January 2021. Using this data, complete COVID-19 and vaccination histories could be deduced for all participants of NIS635. Details for all subcohorts are given in Table 1.

**Table 1:**
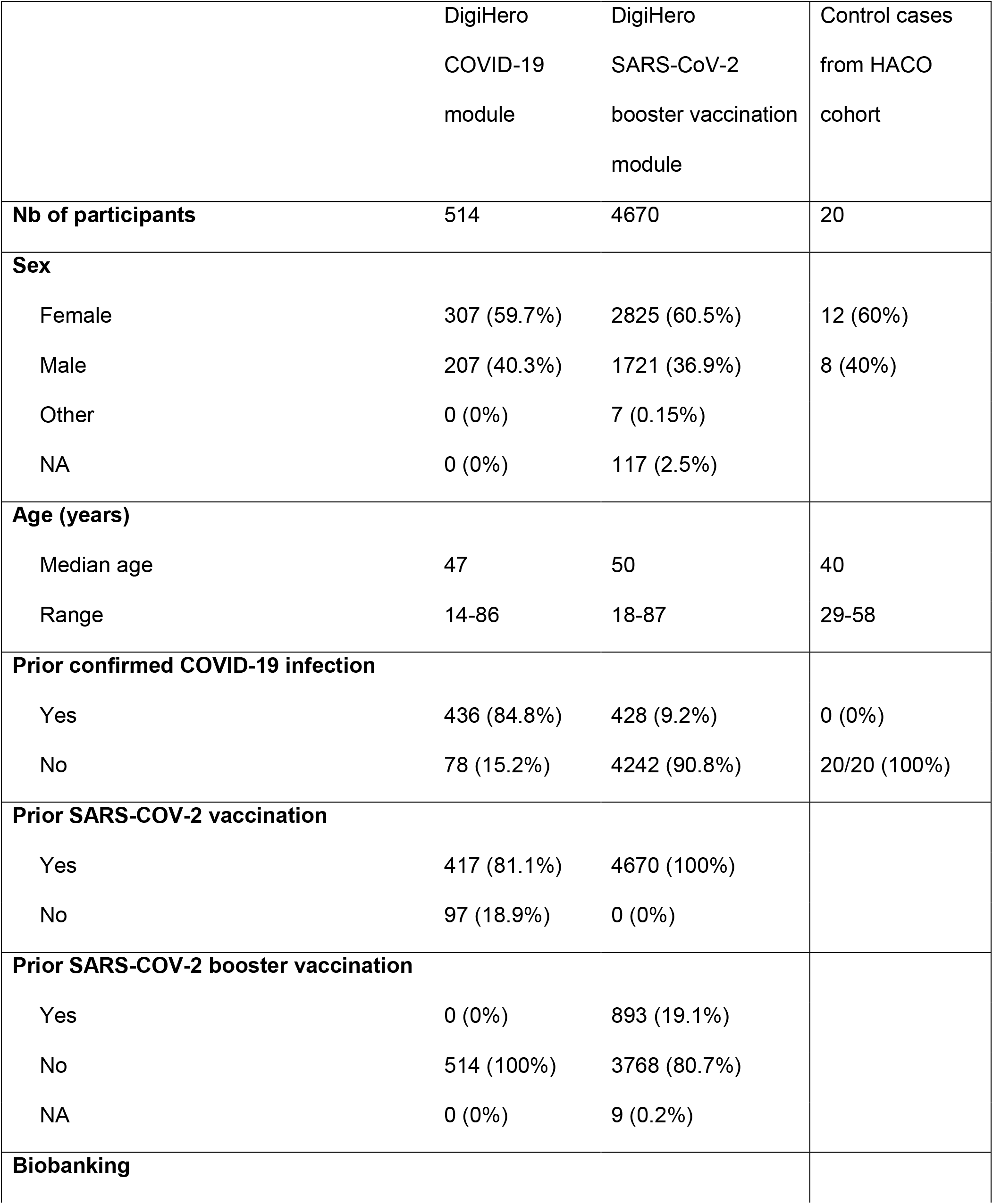
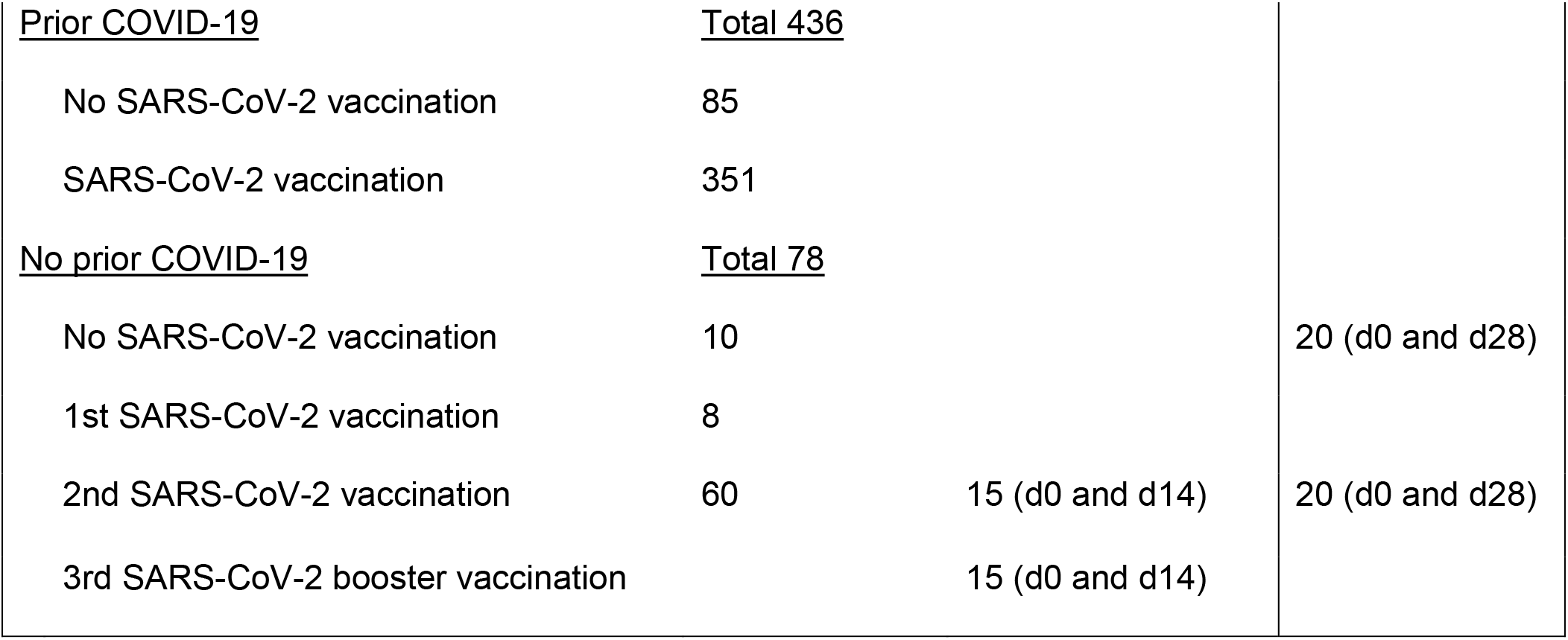
Characteristics of participants in the DigiHero COVID-19 and SARS-CoV-2 booster vaccination modules and the HACO subcohort used for NIS635.

In the DigiHero SARS-CoV-2 “booster” vaccination survey, the majority of participants indicated to have already received their third vaccination or to be planning to receive it shortly (Figure 1A). The time between completion of the primary vaccination series and the third booster vaccination is shown in Figure 1B. The majority of participants received BioNTech/Pfizer as their third vaccine (Figure 1C). The tolerability of the “booster” was roughly comparable to that of first and second SARS-CoV-2 vaccinations (Figure 1D). Most frequent side effects were local reactions, fatigue, and headache with comparable tolerability of both mRNA vaccines (Figure 1E).

**Figure 1:**
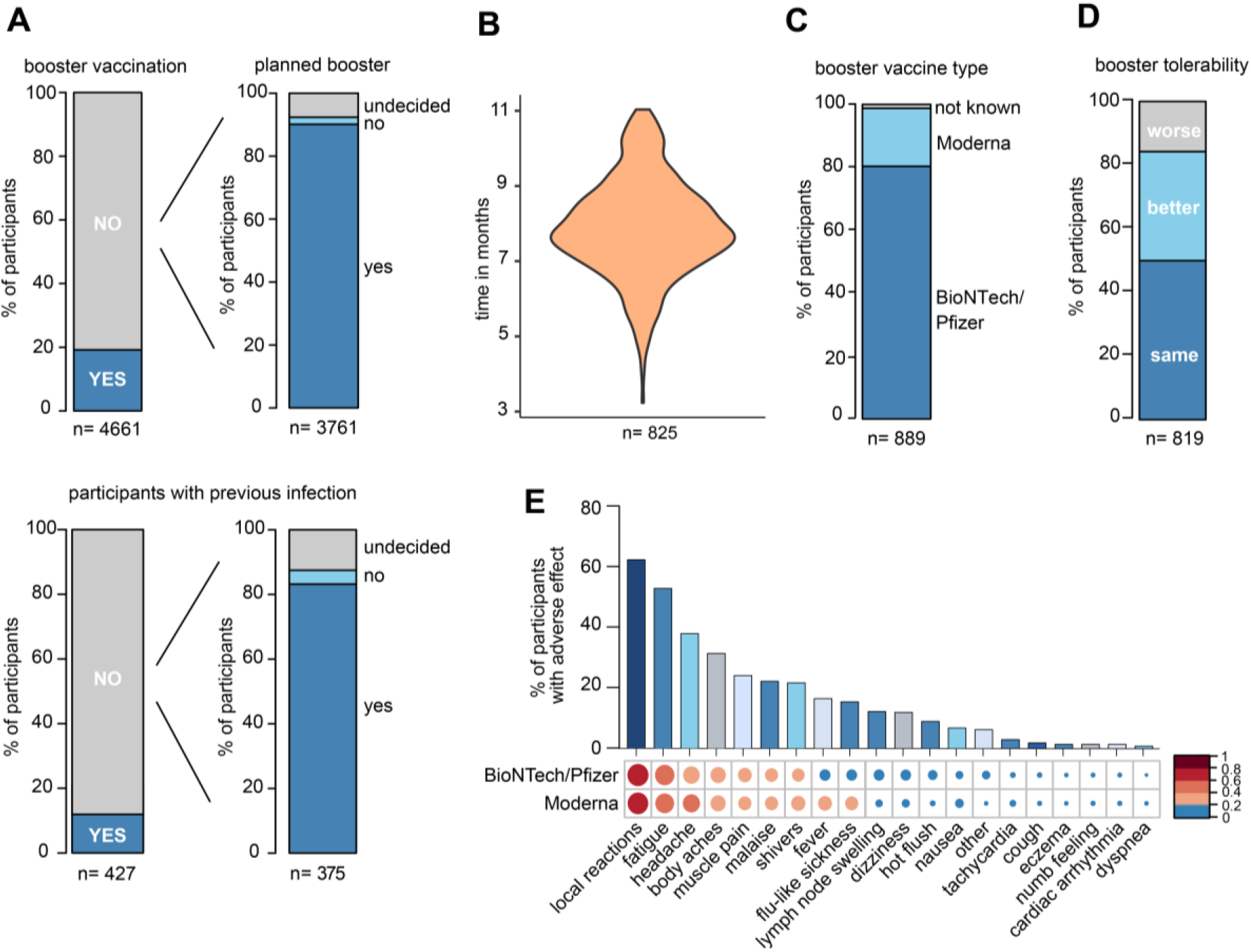
Survey data from the DigiHero SARS-CoV-2 booster vaccination module. (A) Statistics of participants with prior or planned third SARS-CoV-2 vaccination (booster). (B) Time between completion of the primary vaccination series and third vaccination in months. (C) SARS-CoV-2 third vaccination type. (D) Tolerability of the third vaccination compared to previous SARS-CoV-2 vaccinations. (E) Adverse events upon third vaccination.

Based on this survey, 15 participants without prior COVID-19 infection that planned a third vaccination within the next four weeks were randomly chosen and invited to donate blood before and after their third vaccination for antibody and B cell repertoire NGS analyzes.

### SARS-CoV-2 antibodies after infection, priming vaccinations, third vaccination and hybrid immunity

All biobanked samples indicated in Table 1 were tested for S1 and NCP antibodies by ELISA. Figure 2A shows the distribution of antibody levels for the different subgroups. Participants vaccinated after infection (hybrid immunity) and participants after their third “booster” vaccination achieved the highest S1 antibody levels followed by previously uninfected participants that had only completed their primary vaccination series (Figure 2A). Only few individuals showed elevated NCP antibody levels despite having indicated no prior COVID-19 potentially pointing at unrecognized previous infection. All other participants showed antibody levels compatible with their infection/vaccination status. In all participants with matched pre- and post-vaccination samples, clear antibody increases were noted with highest levels after the third vaccination (Figure 2B).

**Figure 2:**
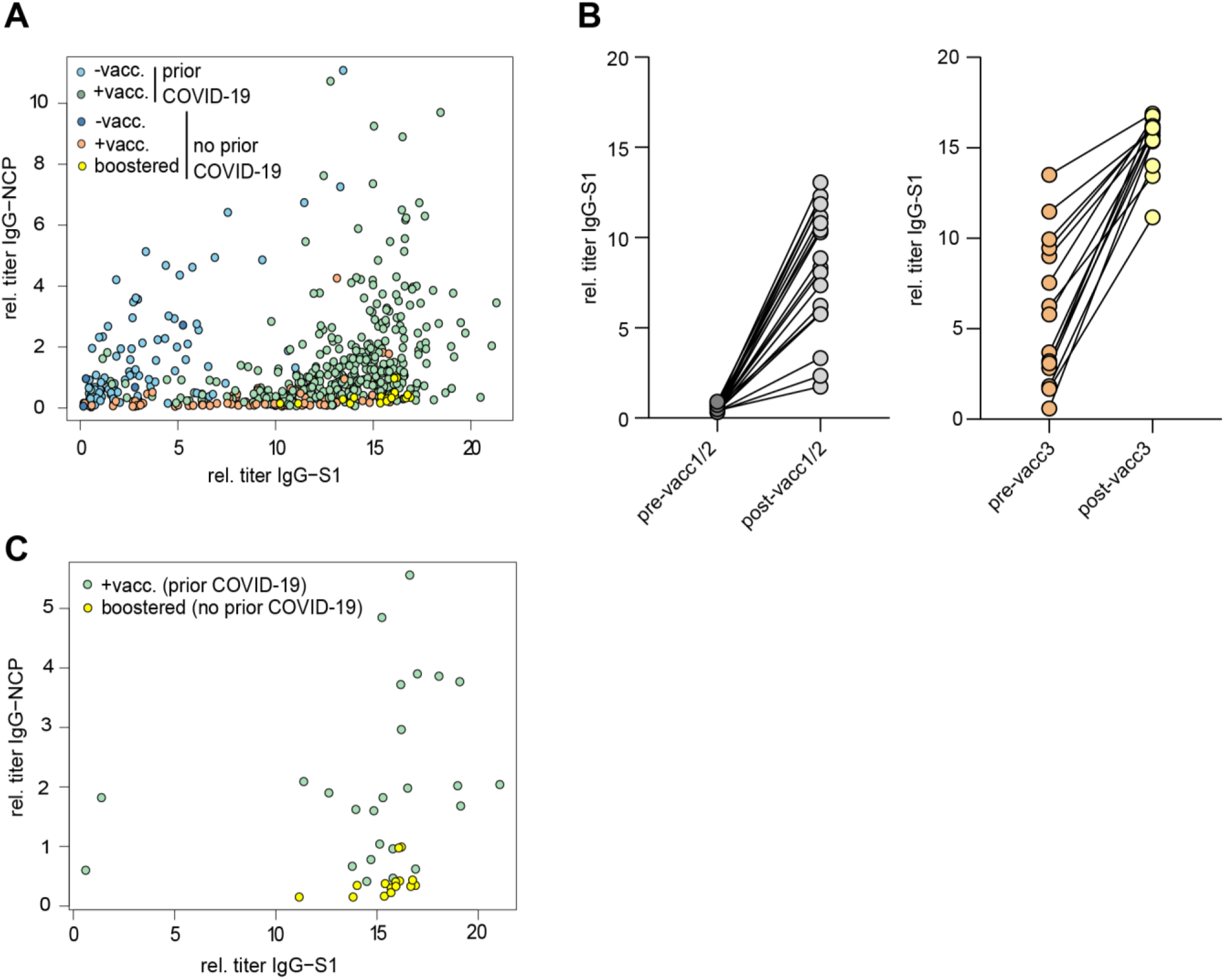
Antibodies against the S1 domain of the spike (S) protein and the nucleocapsid protein (NCP) of SARS-CoV-2. (A) Comparison of IgG-NCP and IgG-S1 antibodies in vaccinated individuals with or without prior COVID-19 infection. (B) Matched IgG-S1 antibody titers of previously uninfected individuals prior to and after the primary vaccination series (pre-/post-vacc1/2) and the third vaccination (pre-/post-vacc3). (C) Comparison of IgG-NCP and IgG-S1 antibodies between previously infected participants that received a subsequent vaccination (green) and previously uninfected participants with three vaccinations (yellow). Both types of blood samples were collected at a maximum of 4 weeks from last vaccination.

Next, we compared antibody levels in participants with hybrid immunity to those after three vaccinations. Given the over-time decay in antibody titers both after infection and vaccination (16, 17, 20, 25–28), we included only participants in this analysis who donated blood in a standard interval of 2-4 weeks after the last vaccination dose. This analysis showed that antibody levels were similarly high in both subsets indicating that the third “booster” dose may mimic the hybrid-like response observed in individuals after infection and vaccination (Figure 2C).

### Global B cell immune metrics after priming and third vaccinations

Matched blood samples prior to and after the first/second or third vaccination were subjected to next-generation sequencing of the B cell receptor repertoire. The time points of blood collection are shown in Figure 3A. All patients included in this analysis had received mRNA vaccines; 8 of 15 participants received the BionTech/Pfizer vaccine as third vaccination. There were no global differences in immune repertoire metrics such as diversity, richness or clonality across groups (Figure 3B). In addition, B cell repertoire somatic hypermutation rates of IGHV genes that reflect antigen-mediated affinity maturation were roughly identical on the global immune repertoire level (Figure 3C). There were, however, trends in repertoire connectivity: Pre-vaccination B cell repertoires showed lowest connectivity between B cell clonotypes while samples taken after the third vaccination showed highest connectivity (Figure 3D, lower part). B cell connectivity plots of two patients with representative connectivity levels are shown in the upper part of Figure 3D.

**Figure 3:**
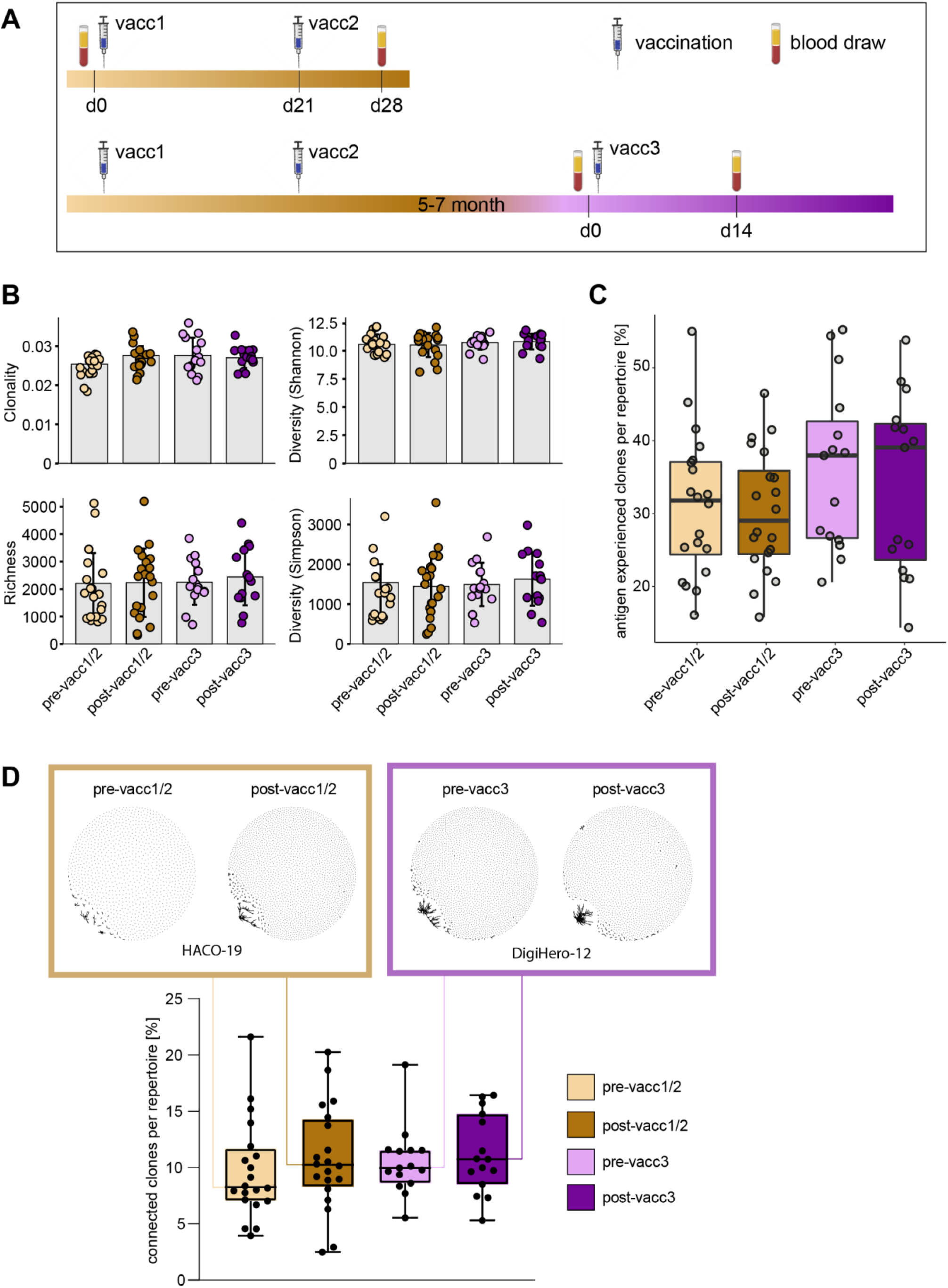
Matched pre- and post-SARS-CoV-2 vaccination blood sampling and global B cell repertoire analysis. (A) Vaccination and blood sampling scheme. (B) Broad B cell repertoire metrics. Bars indicate mean ± standard deviation. (C) Percentage of antigen experienced clones with somatic hypermutation (<98% identity to germline) per B cell repertoire. Box and whiskers plot are shown in the style of Tukey. (D) Quantitative connectivity analysis of B cell clones per repertoire. Boxes outline 25^th^ to 75^th^ percentile with a line at the median and whiskers from minimum to maximum. Petri dish plots of two representative pre-/post vaccination B cell repertoires are shown in brown and violet boxes on top (patients HACO-19 and DigiHero-12).

### SARS-CoV-2 specific B cell clonotypes prior to and after first/second versus third vaccination

While global B cell repertoire metrics were rather stable across the studied subgroups, we hypothesized that the subrepertoire of B cells with known SARS-CoV-2 specificity may provide more insight into affinity maturation in response to vaccination. We, therefore, searched our set of immune repertoires for 3,195 known SARS-CoV-2 antibody sequences (29) to determine blood circulation of B cells carrying SARS-CoV-2 reactive B cell receptors. 1,147 thereof derived from neutralizing SARS-CoV-2 antibodies. Interestingly, blood-circulation of such B cells appeared to be increased shortly after the first two vaccinations (Figure 4A and 4B). After the third vaccination, we also noted increases as compared to the matched time point before the third vaccination (Figure 4A and 4B). Yet, in absolute numbers, the increase in blood circulation of these clones was lower than after the primary vaccination series.

**Figure 4:**
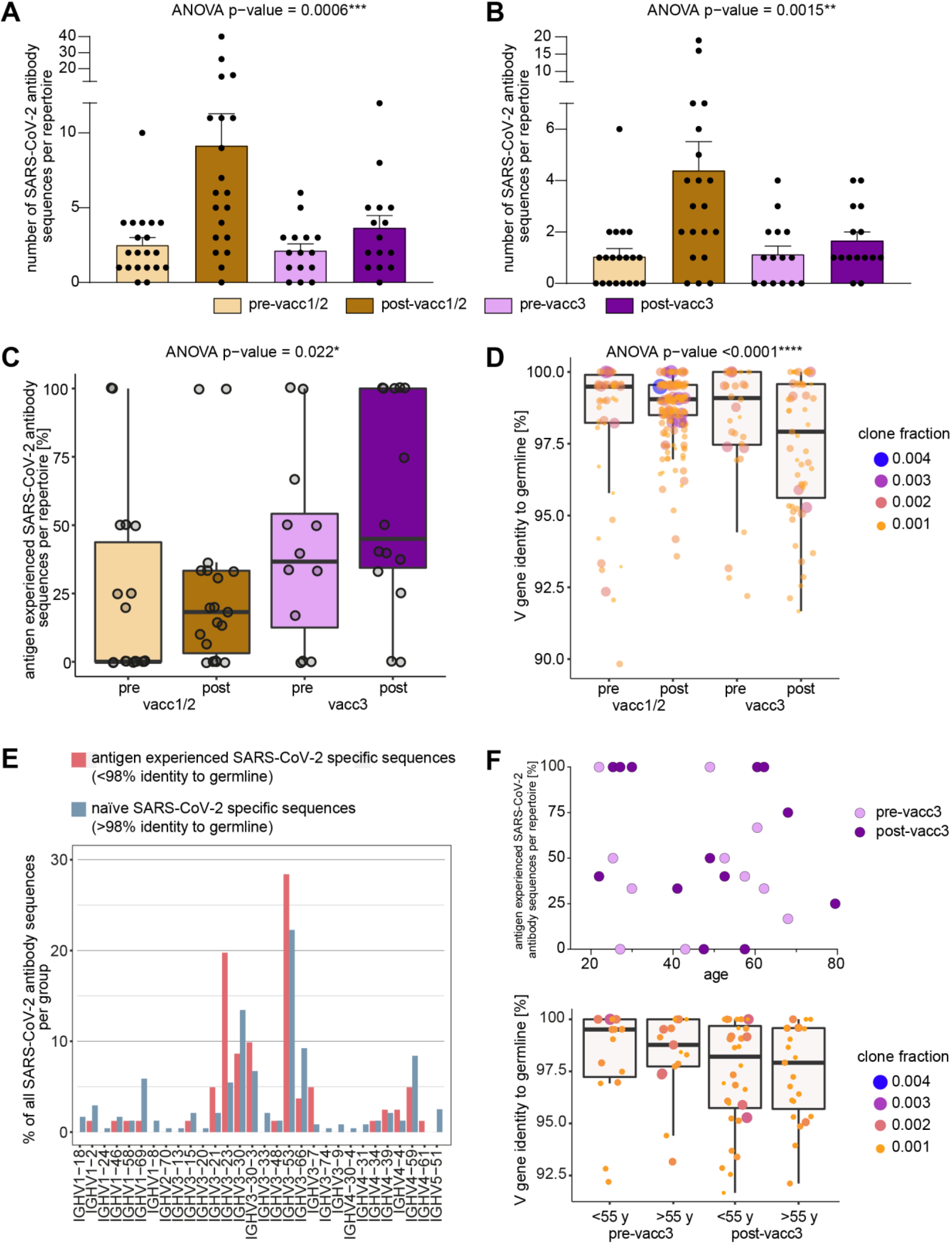
Search of SARS-CoV-2 directed B cell clonotypes in matched pre- and post-vaccination blood samples. Search of 3,195 total (A) and 1,147 neutralizing (B) antibody sequences in all immune repertoires. Bars indicate mean ± s.e.m. (C) Somatic hypermutation analysis of SARS-CoV-2 directed antibody sequences. The percentage of antigen experienced clones within the SARS-CoV-2 specific subrepertoires per patient is shown. (D) Somatic hypermutation load per SARS-CoV-2 directed antibody sequence is shown. Respective clone fractions are coded by color/size. Box and whiskers plot are shown in the style of Tukey. Ordinary one-way ANOVA was performed as statistical test. (E) V gene usage in naïve versus antigen-experienced SARS-CoV-2 directed antibody sequences. (F) Somatic hypermutation analysis of SARS-CoV-2 directed antibody sequences before and after the third vaccination (pre-/post-vacc3) in correlation to participant’s age. The percentage of antigen experienced clones within the SARS-CoV-2 specific subrepertoires per patient is shown in the upper panel. Somatic hypermutation load per SARS-CoV-2 directed antibody sequence is shown before and after the third vaccination (pre-/post-vacc3) subdivided into age groups of </> 55 years (y) in the lower panel. Respective clone fractions are coded by color/size. Box and whiskers plot are shown in the style of Tukey.

In a next step, we determined somatic hypermutation rates of SARS-CoV-2 specific clones from matched pre- and post-vaccination samples. We reasoned that the rate of somatically hypermutated clones should increase with the number of applied vaccinations. Indeed, we found a continuous increase in the fraction of somatically hypermutated B cell clones within the SARS-CoV-2 specific repertoire from pre-vaccination samples to samples acquired after the third vaccination (Figure 4C). Interestingly, the rate of somatically hypermutated SARS-CoV-2 specific clones was lower after completion of the primary vaccination series than prior to the third vaccination. This suggests that even “short-lived” mRNA vaccines trigger affinity maturation of B cells over months in line with recent data (22). Although the somatic hypermutation load per SARS-CoV-2 specific sequence numerically increased in primed participants, it was dramatically boosted by the third vaccination (Figure 4D). This was especially observed for BCR sequences encoded by the IGHV3-21, IGHV3-23, IGHV3-53 and IGHV3-30-3 genes (Figure 4E) which have been already linked to S-reactive antibodies with exceptional neutralizing potency (12, 30–34). The amount of hypermutation did not correlate with age (Figure 4F).

### Maturation trajectories in the primary vaccination series and upon third vaccination

To be able to analyze individual maturation trajectories of B cells in the primary vaccination series versus upon third vaccination, we set out to identify developmental B lineages in all individual participants and to track them across the vaccination period. A B cell lineage is a group of clonotype-defined B cells that share a common V and J gene and a CDR3 sequence differing only in up to 10% of its amino acid positions (35). While we did not detect expanding B cell lineages in a substantial proportion of participants receiving their primary vaccination series, we found expanding lineages in the majority of patients receiving their third vaccination (Figure 5A). The repertoire space taken up by these vaccine-induced expanding B cell lineages was substantially higher after the third vaccination than after completion of the primary vaccination series (Figure 5B). The stream plots in Figure 5C and Supplementary Figure 1 show the development of B cell lineages from the pre-to the post-vaccination time point in all investigated cases. All overlapping, expanding lineages are shown as individual colored stream mapping its frequency within the pre- and post-vaccination repertoire. Detailed analysis of these maturation trajectories showed that the precursors of highly mutated post-booster clones were either naive or antigen-experienced cells that were mobilized into a secondary round of somatic hypermutation, most likely through a second recruitment to a germinal center. Exemplary mutational trajectories induced by the third vaccination starting from naive or antigen-experienced clones are shown in Figure 5D.

**Figure 5:**
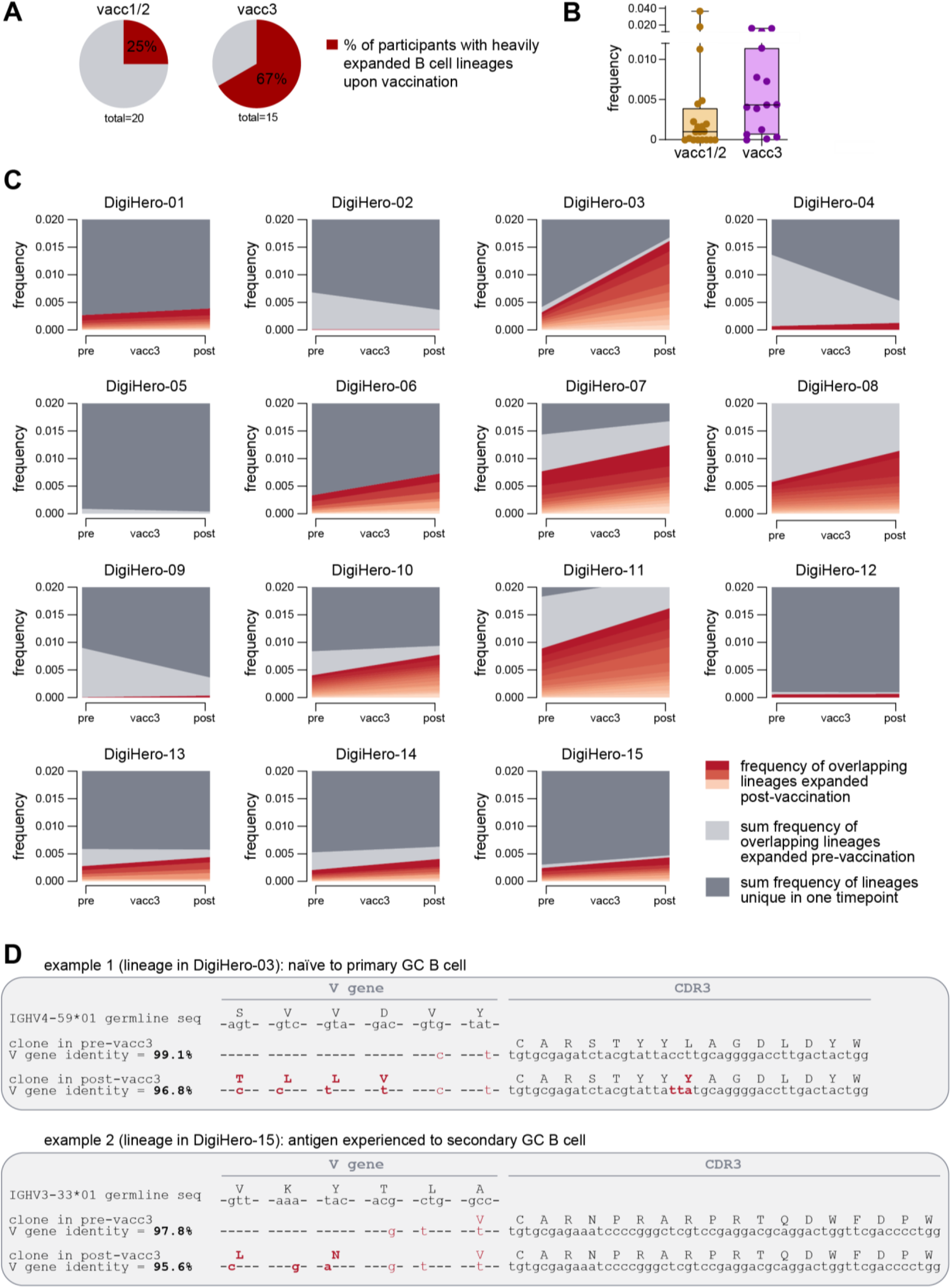
Expanding B cell lineages upon vaccination. B cell lineages in individual patients pre- and post-vaccination were constructed based on V and J gene identity as well as CDR3 sequence homology. (A) The percentage of participants with heavily expanded B cell lineages (more than 0.3% frequency within the post-vaccination repertoire taken up by overlapping expanded lineages) after the first/second (vacc1/2) or third vaccination (vacc3) are shown as pie charts. (B) Repertoire frequency of expanding B cell lineages at the post-vaccination time point. (C) Stream plots showing expanding B cell lineages in patients receiving their third SARS-CoV-2 vaccination. (D) Exemplary detailed somatic hypermutation analysis of two antibody sequences. GC = germinal center. Seq = sequence.

## Discussion

Dissection of infection-, but especially vaccine-induced B cell immunity to SARS-CoV-2 has become even more a priority in light of the advent of SARS-CoV-2 variants of concern that differ from the ancestral strain in transmissibility and immune evasion. In the second half of 2021, when the majority of blood samples for this analysis were collected, Delta (B. 1.617.2) was the dominant SARS-CoV-2 variant worldwide. The Omicron variant B.1.1.529 has been first reported to WHO on the 24^th^ of November 2021 and thereafter spread across the globe at unprecedented rate. In January 2022, Omicron replaced Delta as the dominant variant accompanied by a record of 15 million new COVID-19 cases worldwide in a single week. Both Delta and Omicron variants cause concerns also in fully vaccinated populations since the current vaccines are targeted at the ancestral SARS-CoV-2 strain (36–43). In addition, it was also reported that the type and/or sequence of different exposures triggers SARS-CoV-2-directed immune responses varying in specificity and neutralizing potency (44). It is therefore crucial to understand the biology of SARS-CoV-2 vaccination in detail to design better vaccination protocols and eventually vaccine updates providing sufficient protection against emerging variants.

In the study presented here, we show complex B cell maturation trajectories induced by the third “booster” vaccination that by far exceeded the level of somatic hypermutation measurable directly after completion of the primary vaccination series. Interestingly, the strong mutational activity was essentially restricted to few lineages that were present and mutated already before the third vaccination and therefore likely involved in previously induced memory B cells that were once again recruited to the lymph node’s germinal center for further refinement. Notably, we observed high mutational rates especially in IGHV3-21, IGHV3-23, IGHV3-53 and IGHV3-30-3 genes that were linked to antibodies isolated from elite neutralizers that have previously shown neutralizing potency against variants of concern after infection with the ancestral strain (12, 30–34). Interestingly, we did not observe any age-restriction for this maturation process indication that also older people benefit from a third vaccination although this needs further validation due to the limited size of the analyzed cohort. Since somatic hypermutation reflects affinity maturation, our data is not only well compatible with the strong increase in neutralization potential towards the ancestral strain induced by the third vaccination (45), it also may explain why individuals after a recent third vaccination are usually protected from infection with the Delta variant that shows only few immune-evasive S protein mutations (46–48). Since high affinity towards the ancestral strain is a prerequisite for variant neutralization, our data may also explain why the third vaccination but not the primary vaccination series produces some level of protection against infection with the Omicron strain that harbors a large number of immune-evasive mutations within the S protein (49, 50).

We found a gap between the fraction of somatically hypermutated SARS-CoV-2 directed B cell clones after completion of priming and directly before boosting. This suggested that SARS-CoV-2 vaccines might trigger long-term maturation and affinity selection in the B cell compartment over months even with mRNA vaccines that consist in short-lived injected molecules. This data is well compatible with recent findings from Sokal et al. (24) that show continuous maturation of B cells over months after vaccination by flow cytometry. From a translational perspective, this data may suggest that the shortening of the interval between priming and boosting may come at a price of lower booster efficacy if the time span for ongoing B cell maturation and affinity selection is too short. Increasing the time interval between the first and the second dose has been shown to promote effectiveness in the AstraZeneca vector vaccine trial for the first two vaccinations (51). At the same time, this needs to be balanced against the current risk associated with the inability of a primary vaccination series to protect against the current Omicron variant.

The third vaccination rapidly generated a considerable somatic hypermutation load in SARS-CoV-2 specific B cell receptor clonotypes given that the time interval between the pre- and post-booster vaccination sampling was only 14 days. To achieve a somatic hypermutation rate of the IGHV gene between 2 and 10% (typical for the antigen-experienced B cells seen in our study), between 6 and 30 nucleotide exchanges are required per clone. Given the increased somatic hypermutation rate of immunoglobulin genes of approximately 10(−3) mutations per base pair per cell division (52), more than three cell divisions are necessary to statistically exchange one nucleotide per IGHV gene. This very rough calculation may show the high proliferative stimulus of the third booster vaccination. The clinical correlate thereof may be the rather high rate of ipsilateral axillary lymph node swelling compared to other cohorts (53) observed by almost 10% of our DigiHero participants after the third vaccination.

Taken together, our data show that the primary vaccination series quickly generates antibodies from B cells that have only undergone low-level affinity maturation and may therefore not be protective for immune-escape viral variants such as Omicron B.1.1.529. Our analyzes confirm the role of the third SARS-CoV-2 “booster” to generate affinity-matured clones and mobilize them for antibody production.

## Methods

### Non-interventional study design and biobanking of NIS635

This study was registered as non-interventional study (NIS) at the Paul-Ehrlich-Institute (NIS635). It consisted of data and biological samples collected in the DigiHero and HACO cohorts.

DigiHero is a population-based cohort study for digital health research in Germany conducted in the city of Halle (Saale) which registered 8,077 participants until November 2021. The recruitment was conducted in different waves and included mailed invitation to all 129,733 households in Halle as well as promotion via media. The study was approved by the institutional review board (approval number 2020-076). Its digital design allowed targeted invitation of participants to modules that included different surveys and blood biobanking subprojects. The COVID-19 module of DigiHero recruited participants with prior positive SARS-CoV-2 testing in their households. Until December 2021, 514 individuals had completed the survey on their COVID-19 history as well as on their vaccination status and had donated blood for this module at this data cut. These samples were used for antibody analyzes. The SARS-CoV-2 booster vaccination module of DigiHero recruited participants willing to respond to a survey on the third SARS-CoV-2 booster vaccination. Until December 2021, 4,670 participants had completed the survey. Fifteen randomly chosen participants without prior COVID-19 infection donated blood prior to and on day 14 (d14) after booster vaccination at this data cut. These samples were used for antibody and B cell repertoire next-generation sequencing analyzes.

As a reference within NIS635, 40 samples from 20 uninfected control cases completing their primary vaccination series with the BioNTech/Pfizer vaccine were recovered from the biobank of the Halle COVID-19 cohort (HACO). These 20 control cases had donated blood prior to and at d28 of their primary vaccination series (d7 after the second vaccination).

Informed written consent was obtained and the study was approved by the institutional review board (approval number 2020-039). These samples were used for antibody and B cell repertoire next-generation sequencing analyzes.

Table 1 summarizes all relevant participant numbers, their basic characteristics and biological samples used in NIS635. The study was conducted in accordance with the ethical principles stated by the Declaration of Helsinki. Informed written consent was obtained from all participants or legal representatives.

### Sample collection

The collected plasma samples were isolated by centrifugation of whole blood for 15 min at 2,000xg, followed by centrifugation at 12,000xg for 10 min and stored at −80°C. Peripheral mononuclear cells (PBMC) were isolated by standard Ficoll gradient centrifugation. Genomic DNA was extracted from PBMCs using the GenElute Mammalian Genomic DNA Miniprep Kit (Sigma-Aldrich, St. Louis, USA).

### SARS-CoV-2 antibody profiling

Antibodies against the S1 domain of the spike (S) protein and the nucleocapsid protein (NCP) of SARS-CoV-2 were determined by Anti-SARS-CoV-2-ELISA IgA/IgG and Anti-SARS-CoV-2-NCP-ELISA kits from Euroimmun (Lübeck, Germany). Readouts were performed at 450 nm using a Tecan Spectrophotometer SpectraFluor Plus (Tecan Group Ltd., Männedorf, Switzerland).

### Next-generation sequencing (NGS) of B cell immune repertoires

Immunosequencing of B cell repertoires was performed as described in (54). In brief, V(D)J rearranged IGH loci were amplified from 500 ng of genomic DNA using a multiplex PCR, pooled at 4 nM and quality-assessed on a 2100 Bioanalyzer (Agilent Technologies). Sequencing was performed on an Illumina MiSeq (paired-end, 2 x 301-cycles, v3 chemistry). Rearranged IGH loci were annotated using MiXCR v3.0.13 (55) and the IMGT 202011-3.sv6 IGH library as reference. Non-productive reads and sequences with less than 2 counts were discarded. All repertoires were normalized to 30,000 reads. Each unique complementarity-determining region 3 (CDR3) nucleotide sequence was considered a clone. Broad repertoire metrics (clonality, diversity, richness) were analyzed as previously described (56). IGHV genes were regarded as somatically hypermutated if they showed < 98% identity to the germline sequence.

### B cell clonotype search algorithm

We searched our IGH repertoires for validated SARS-CoV-2 antibody rearrangements with identical or highly similar CDR3 amino acid sequence (Levenshtein distance of ≤ 2) and identical IGHV-J gene usage as described in (13). The validated SARS-CoV-2 antibody sequences were derived from CoV-AbDab accessed at 17^th^ December 2021 (29) and classified into 3,195 total SARS-CoV-2 binding sequences and 1,147 SARS-CoV-2 neutralizing sequences. A list of the target sequences is provided in Supplementary Table 1.

### B cell network analysis

To calculate network connectivity in BCR repertoires, we used the Levenshtein distance of all unique CDR3 amino acid sequences per repertoire using the imnet tool (https://github.com/rokroskar/imnet). Sequences with Levenshtein distance ≤ 3 were connected. For visualization as petri dish plots we used R package igraph and the fruchterman-reingold layout (57). Data analysis and plotting was performed using R version (v4.1.2).

### Identification of B cell lineages

We identified overlapping B cell lineages in the pre- and post-vaccination time point per patient. A B cell lineage was defined as a group of B cell clonotypes that share a common V and J gene and a CDR3 sequence differing only in up to 10% of its amino acid positions (35). Lineages and their evolution were visualised as stream plots with function plot.stacked from (https://www.r-bloggers.com/2013/12/data-mountains-and-streams-stacked-area-plots-in-r/). Data analysis and plotting was performed using R version (v4.1.2).

### Data availability

The herein reported sequence data set has been deposited at the European Nucleotide Archive (ENA) under the accession number PRJEB50803.

### Statistics

Differences between the four groups were analyzed by ordinary one-way ANOVA. Differences between two groups were studied by unpaired, two-tailed student’s t-test. All statistical analyses were performed using R version 4.1.2 and GraphPad Prism 8.3.1 (GraphPad Software, La Jolla, CA, USA).

### Study approval

This study was registered as non-interventional study (NIS) at the Paul-Ehrlich-Institute (NIS635). DigiHero and HACO were approved by the institutional review board (approval numbers 2020-076 and 2020-039). Written informed consent was received prior to participation.

## Author contributions

Idea & design of research project: MB, LP, RM, CG. Supply of critical material (e.g. patient material, cohorts): RM, DS, MB, MG, MGi, SD, BK; Establishment of Methods: LP, CS, EW; Experimental work: LP, CS; Data analysis and interpretation: MB, LP, CS; Drafting of manuscript: MB, LP.

## Acknowledgments

We thank the participants of our DigiHero and HACO cohorts for their great support. Moreover, we thank Christoph Wosiek, Aline Patzschke, Jenny Wehde and Katrin Nerger for excellent technical assistance. This project was partially funded by the CRC 841 of the German Research Foundation (to MB) as well as by the Martin-Luther-University Halle (Saale).

## Supplemental Information

**Supplemental Figure 1: Stream plots showing expanding B cell lineages in patients receiving their primary vaccination series against SARS-CoV-2.** B cell lineages in individual patients pre- and post-vaccination were constructed based on V and J gene identity as well as CDR3 sequence homology.

**Supplemental Table 1: List of SARS-CoV-2 directed sequences used in our search algorithm**.

