## Supplemental Figure 1 for "Rapid hypermutation B cell trajectory recruits previously primed B cells upon third SARS-CoV-2 mRNA vaccination"

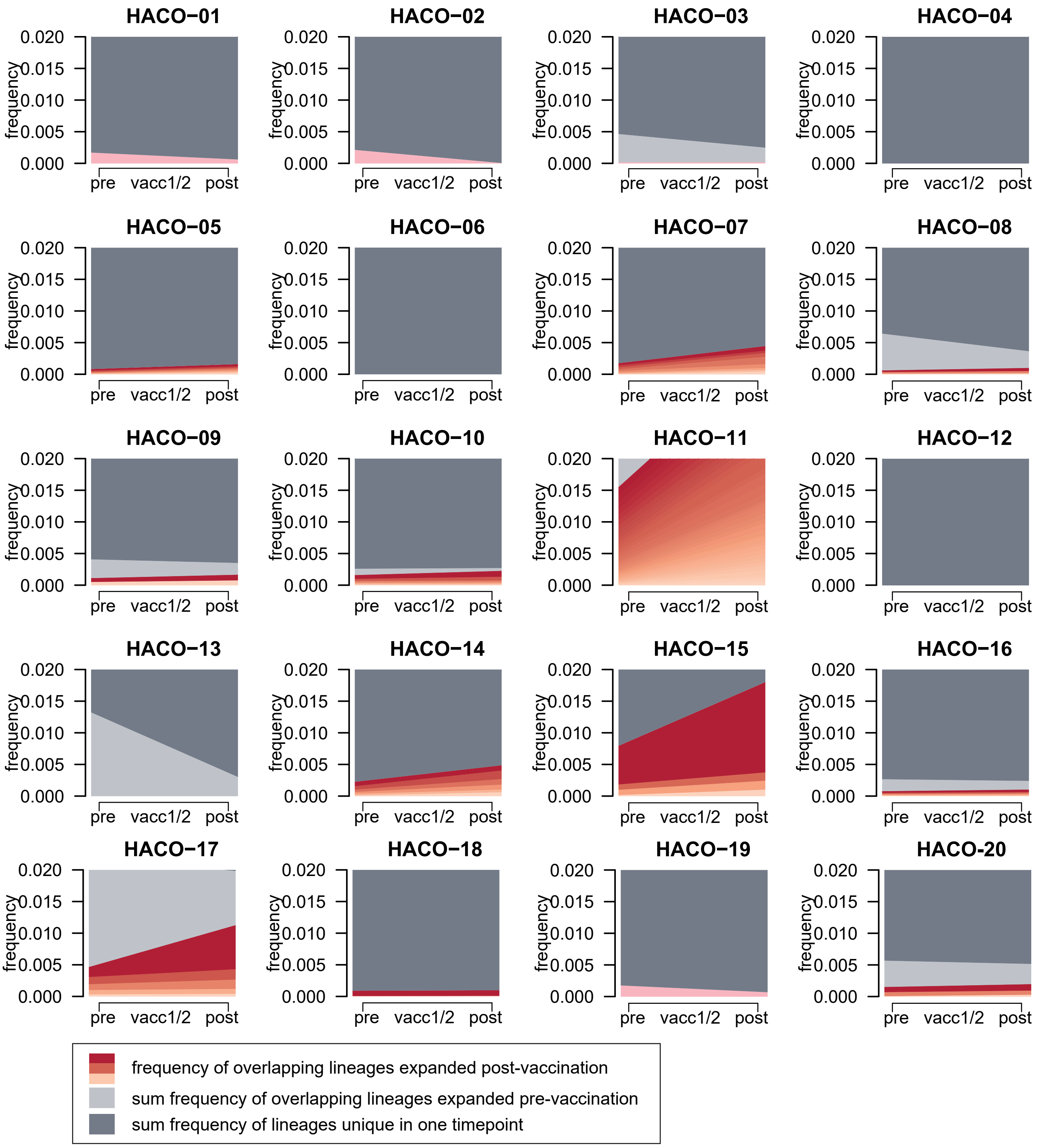


**Supplemental Figure 1: Stream plots showing expanding B cell lineages in patients receiving their primary vaccination series against SARS-CoV-2.** B cell lineages in individual patients pre- and post-vaccination were constructed based on V and J gene identity as well as CDR3 sequence homology.
